## Supplementary Figures for "SPOROCYTELESS/NOZZLE acts together with MADS-domain transcription factors to regulate an auxin-dependent network controlling the Megaspore Mother Cell development"

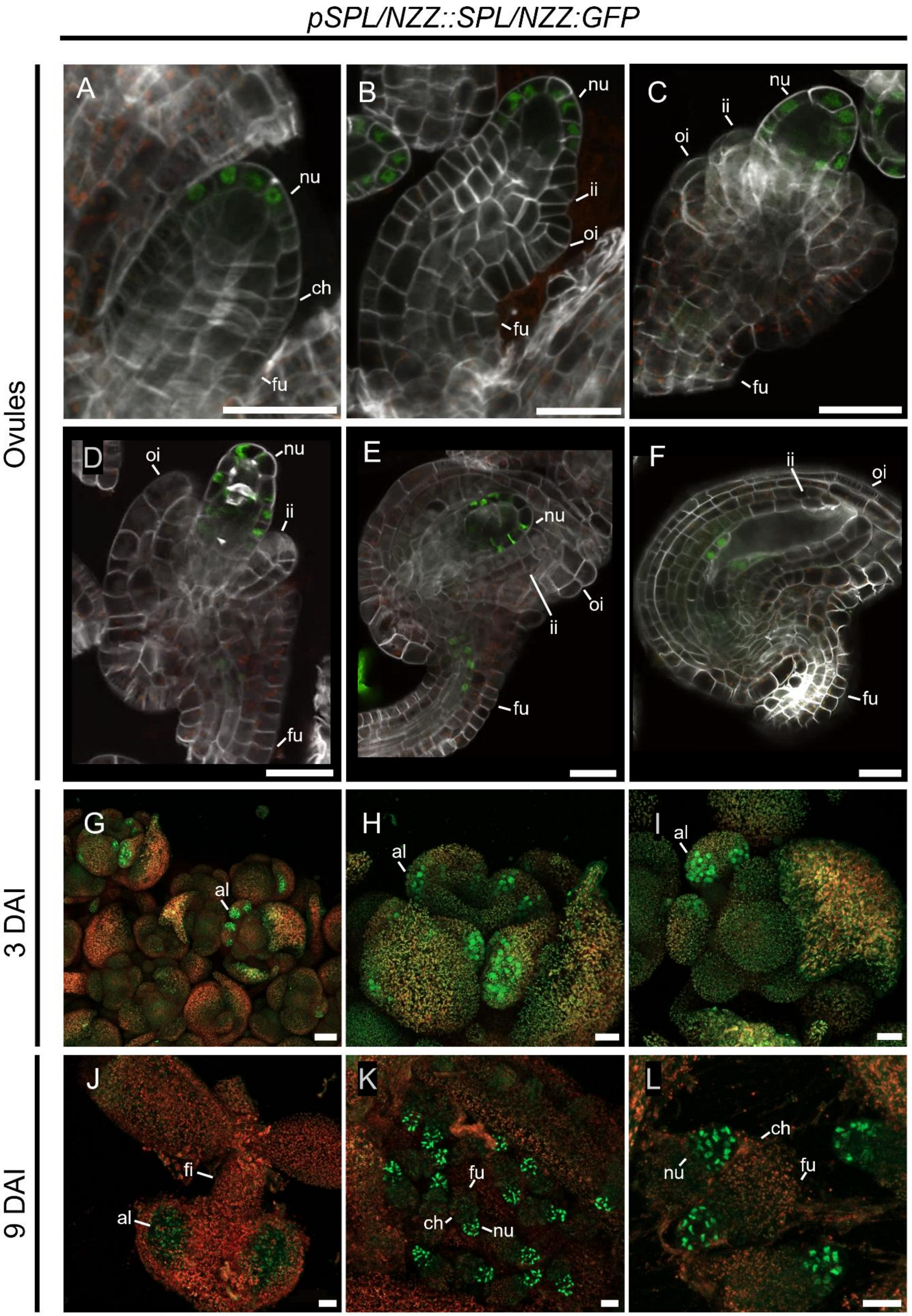


**Supplementary Figure 1: *pSPL/NZZ::SPL/NZZ:GFP* expression.** (**A-F**) *pSPL/NZZ::SPL/NZZ:GFP* expression during different stages of ovule development. SPL/NZZ is localised in the apical L1 cells at stages 1-II (**A**) and 2-II (**B**). At stage 2-

IV (**C**) and 2-V (**D**), SPL/NZZ accumulates in whole nucellar L1 layer. At stage 3-I (**E**), SPL/NZZ is present in the L1 layer and in the central part of the funiculus. At ovule maturity (**F**), the L1 layer degenerates and SPL/NZZ accumulates in the remaining cells. (**G-I**) *pSPL/NZZ::SPL/NZZ:GFP* expression in *pAP1::AP1:GR ap1cal* inflorescences at 3 DAI. At 3 DAI, SPL/NZZ-GFP signal is visible only in developing anthers. (**J-L**) *pSPL/NZZ:SPL/NZZ:GFP* expression in *pAP1::AP1:GR ap1cal* inflorescences at 9 DAI. At 9 DAI, SPL/NZZ-GFP is not visible in anthers anymore (**J**). By contrast, SPL/NZZ-GFP strongly accumulates in ovules nucellus (**K, L**). Abbreviations: nu= nucella; ch= chalaza; fu= funiculus; ii= inner integument; oi= outer integument; al= anther lobe; fi= filament. (**A-F, H-L**) Scale bar = 20 µm. (**G**) Scale bar = 50 µm.

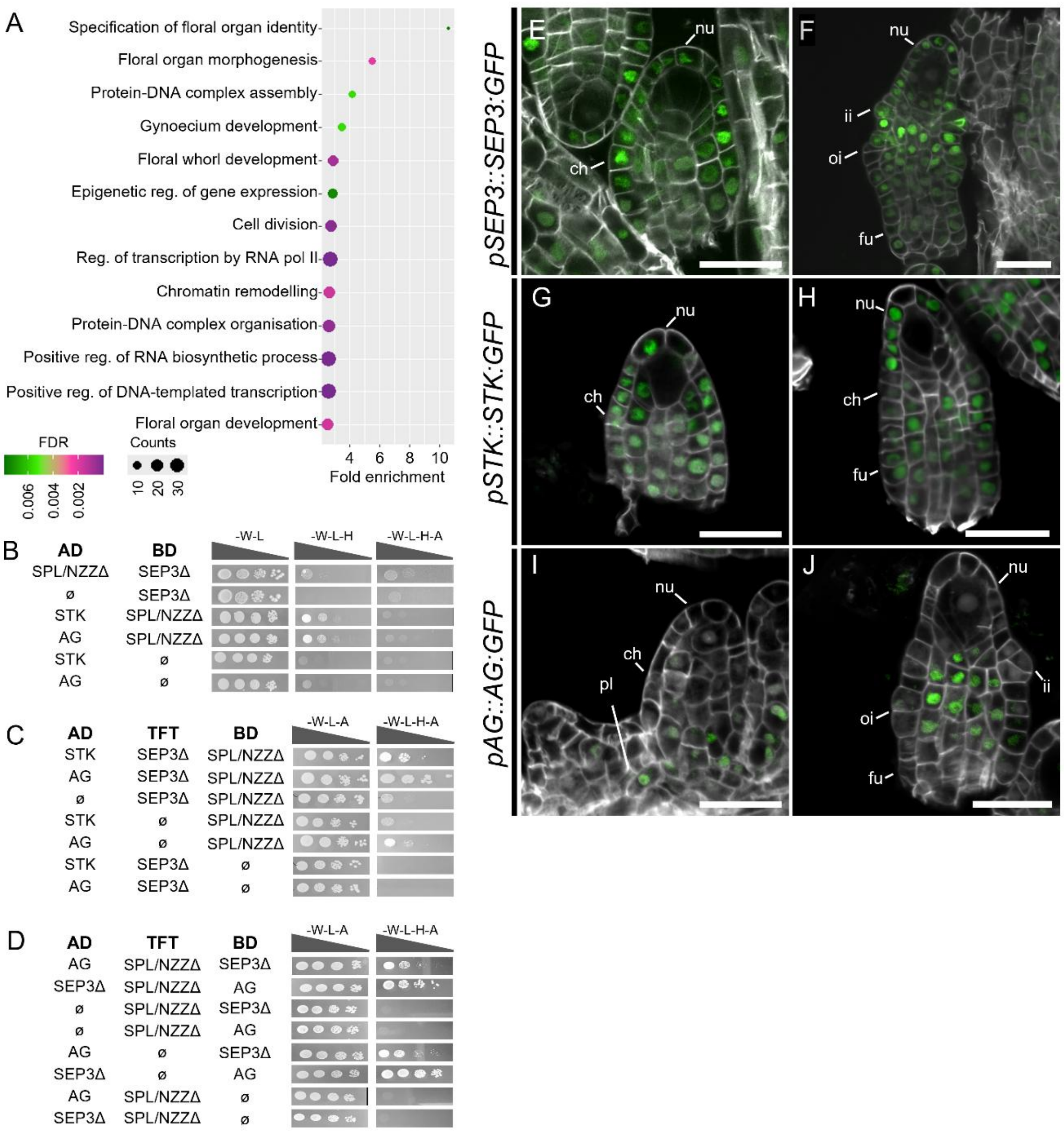


**Supplementary Figure 2: GO analysis of SPL/NZZ interactors identified by the Co-IP/MS, results of yeast-two- and three-hybrid assays and SEP3, STK and AG accumulation patterns in the ovule.** (**A**) Biological process GO terms enrichment analysis of the SPL/NZZ putative direct interactors identified by Co-IP/MS. Enriched Go terms with an FDR ≤ 0.05 were considered significant. (**B**) Yeast-two-hybrid assay showing interaction between SPL/NZZ and MADS-domain TFs (SEP3, STK, AG). The controls are shown in the Figure 1C. Interactions are tested on media depleted of histidine (-W -L -H), or histidine and adenine (-W -L -H -A). For each interaction on the different media, yeast has been spotted at four different concentrations according to a serial dilution (OD 0,5; 1:10; 1:100; 1:1000). (**C, D**) Yeast-three-hybrid assay showing the formation of SPL/NZZ, STK and SEP3 complex, by fusing SPL/NZZ either to the GAL4 BD (**C**) or to the TFT (**D**)**.** Interactions are tested on a medium depleted of histidine (-W -L -H -A). For each interaction, yeast has been spotted at four different concentrations according to a serial dilution (OD 0,5; 1:10; 1:100; 1:1000). (**E, F**) *pSEP3::SEP3:GFP* reporter line showing SEP3 accumulation in ovules at stage 2-I (**D**) and 2-III (**E**). (**G, H**) *pSTK::STK:GFP* reporter line in ovules at stage 2-I (**G**) and 2-II (**H**)**.** (**I, J**) *pAG::AG:GFP* reporter line in ovules at stage 2-I (**I**) and 2-III (**J**)**.** Abbreviations: nu= nucella; ch= chalaza; fu= funiculus; ii= inner integument; oi= outer integument; pl= placenta. Scale bar = 20 µm.

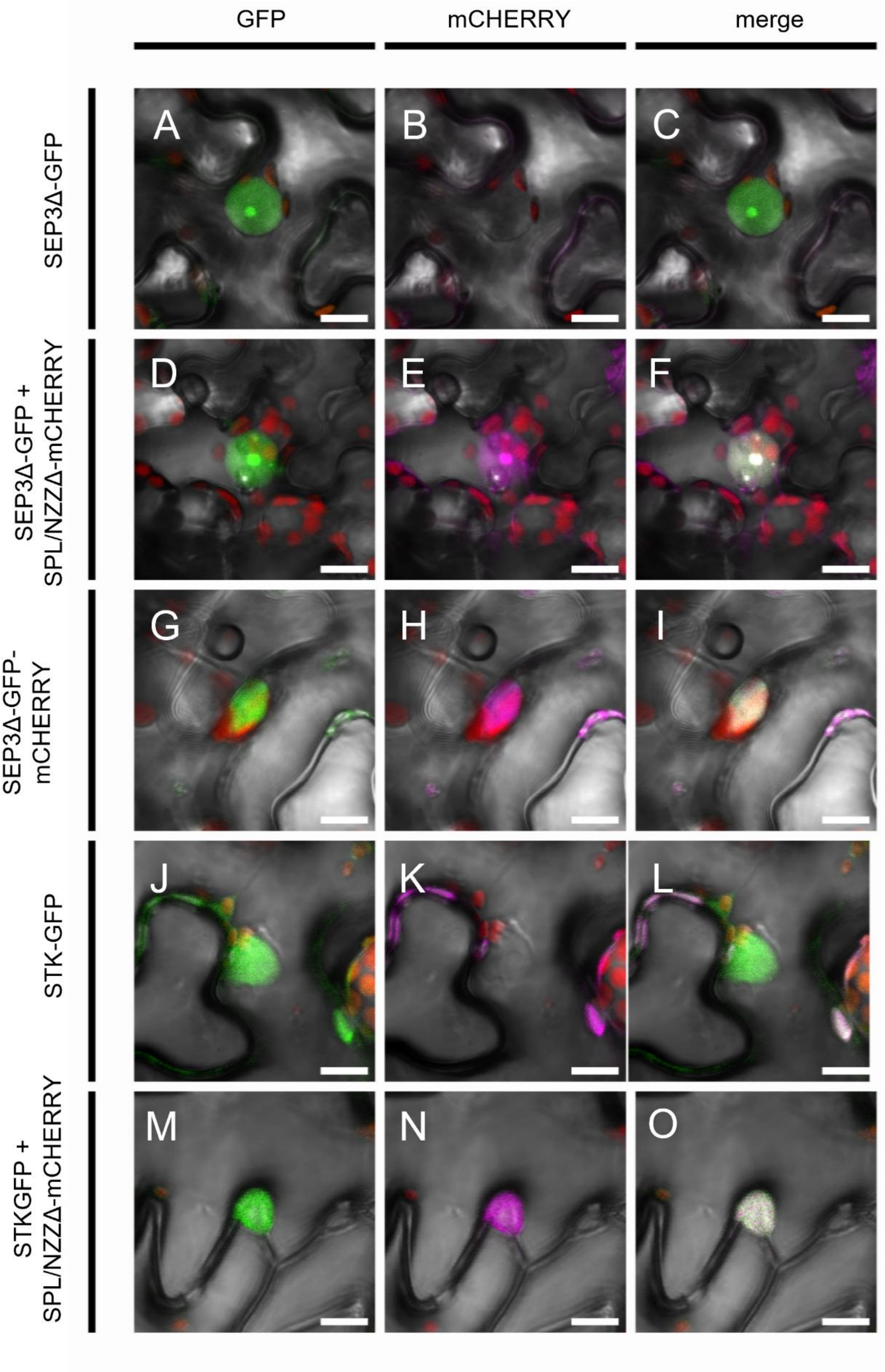


**Supplementary Figure 3: Single and multi-channel images showing GFP and mCHERRY signals in the nuclei represented in Figure 1D.** (**A-C**) GFP channel (**A**), mCHERRY channel (**B**) and merge image of GFP and mCHERRY channels (**C**) of a nucleus expressing SEP3Δ-GFP. (**D-F**) GFP channel (**D**), mCHERRY channel

(**E**) and merge image of GFP and mCHERRY channels (**F**) of a nucleus expressing SEP3Δ-GFP + SPLΔ-mCHERRY. (**G-I**) GFP channel (**G**), mCHERRY channel (**H**) and merge image of GFP and mCHERRY channels (**I**) of a nucleus expressing SEP3Δ-GFP-mCHERRY. (**J-L**) GFP channel (**J**), mCHERRY channel (**K**) and merge image of GFP and mCHERRY channels (**L**) of a nucleus expressing STK-GFP. (**M-O**) GFP channel (**M**), mCHERRY channel (**N**) and merge image of GFP and mCHERRY channels (**O**) of a nucleus expressing STK-GFP + SPLΔ-mCHERRY. Scale Bars: 10 μm.

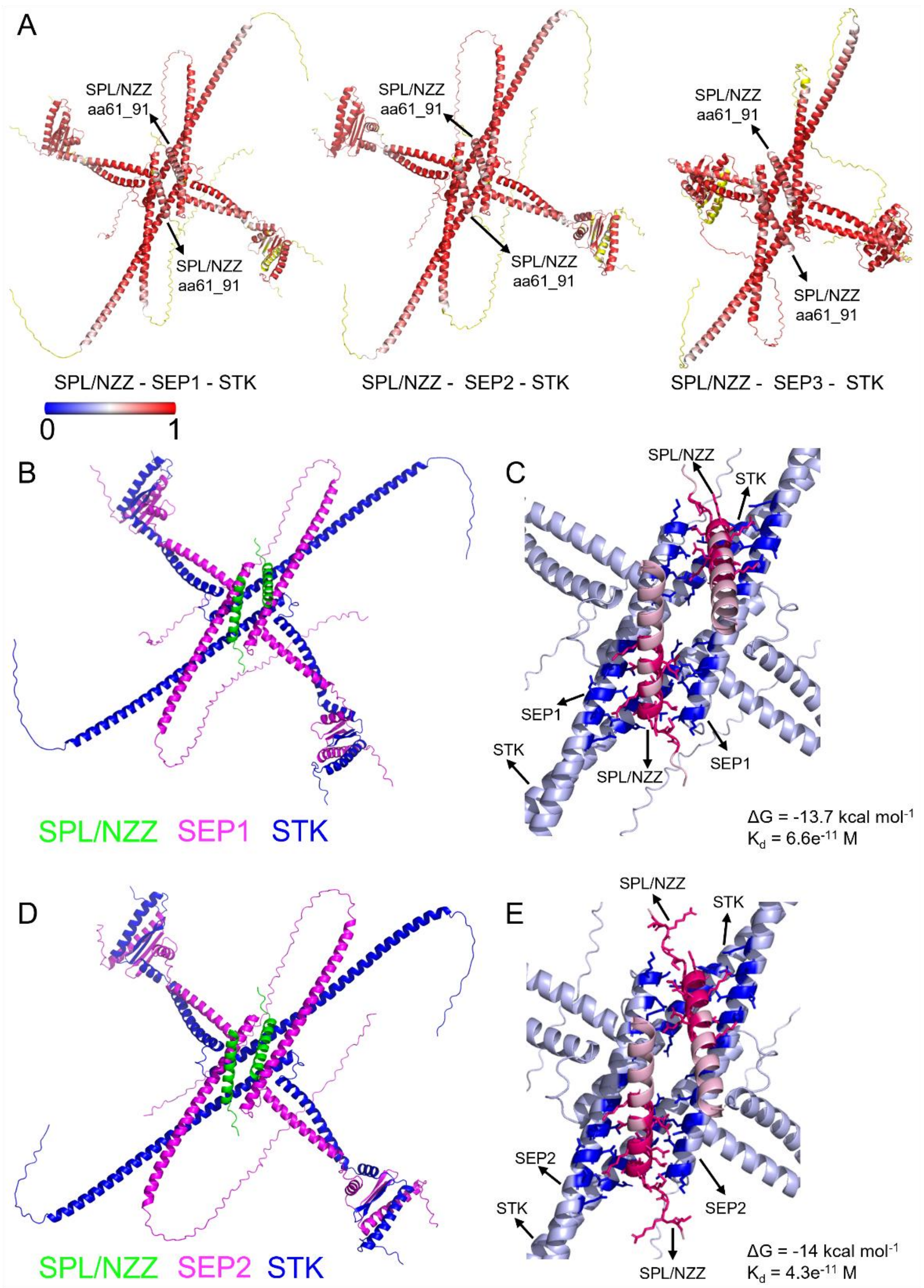


**Supplementary Figure 4: AplhaFold3 predictions of SPL/NZZ-SEPs-STK complexes.** (**A**) SPL/NZZ-SEPs-STK complex predictions coloured according to the prediction confidence. The spectrum bar represents the confidence colours: red colours represent a high confidence (1), while blue colours a low confidence (0). From left to right, SPL/NZZ-SEP1-STK complex, SPL/NZZ-SEP2-STK complex and SPL/NZZ-SEP3-STK complex. (**B**) Prediction of the complex made by two copies of STK (in blue), two copies of SPL/NZZaa61-91 (in green) and two copies of SEP1 (in magenta). (**C**) Close-up view of the SPL/NZZ-SEP1-STK complex according to a binding-affinity prediction. The coloured portion on the complex contains 88 intermolecular contacts between SPL/NZZ and the STK-SEP1 at a maximum distance of 5.5 Å. Particularly, STK and SEP1 amino acids that take contact with SPL/NZZ are showed in blue. SPL/NZZ amino acids that participate in binding to STK and SEP1 are depicted in red. Light blue and light red colouring marks regions where there is no binding. It is possible to notice that one SPL/NZZ molecule interacts with the K-domain of one STK molecule and two SEP1 molecules. The Gibbs free energy (ΔG) for this complex is -13.7 kcal $mol^{-1}$ and the dissociation constant ($K_d$) is 6.6 $e^{-11}$ M. (**D**) prediction of the complex made by two copies of STK (in blue), two copies of SPL/NZZaa61-91 (in green) and two copies of SEP2 (in magenta). (**E**) Close-up view of the SPL/NZZ-SEP2-STK complex according to a binding-affinity prediction. The coloured portion on the complex contains 98 intermolecular contacts between SPL/NZZ and the STK-SEP2 at a maximum distance of 5.5 Å. Particularly, STK and SEP2 amino acids that take contact with SPL/NZZ are showed in blue. SPL/NZZ amino acids that participate in binding to STK and SEP2 are depicted in red. Light blue and light red colouring marks regions where there is no binding. It is possible to notice that one SPL/NZZ molecule interacts with the K-domain of one STK molecule and two SEP2 molecules. The Gibbs free energy (ΔG) for this complex is -14.0 kcal $mol^{-1}$ and the dissociation constant ($K_d$) is 4.3 $e^{-11}$ M.

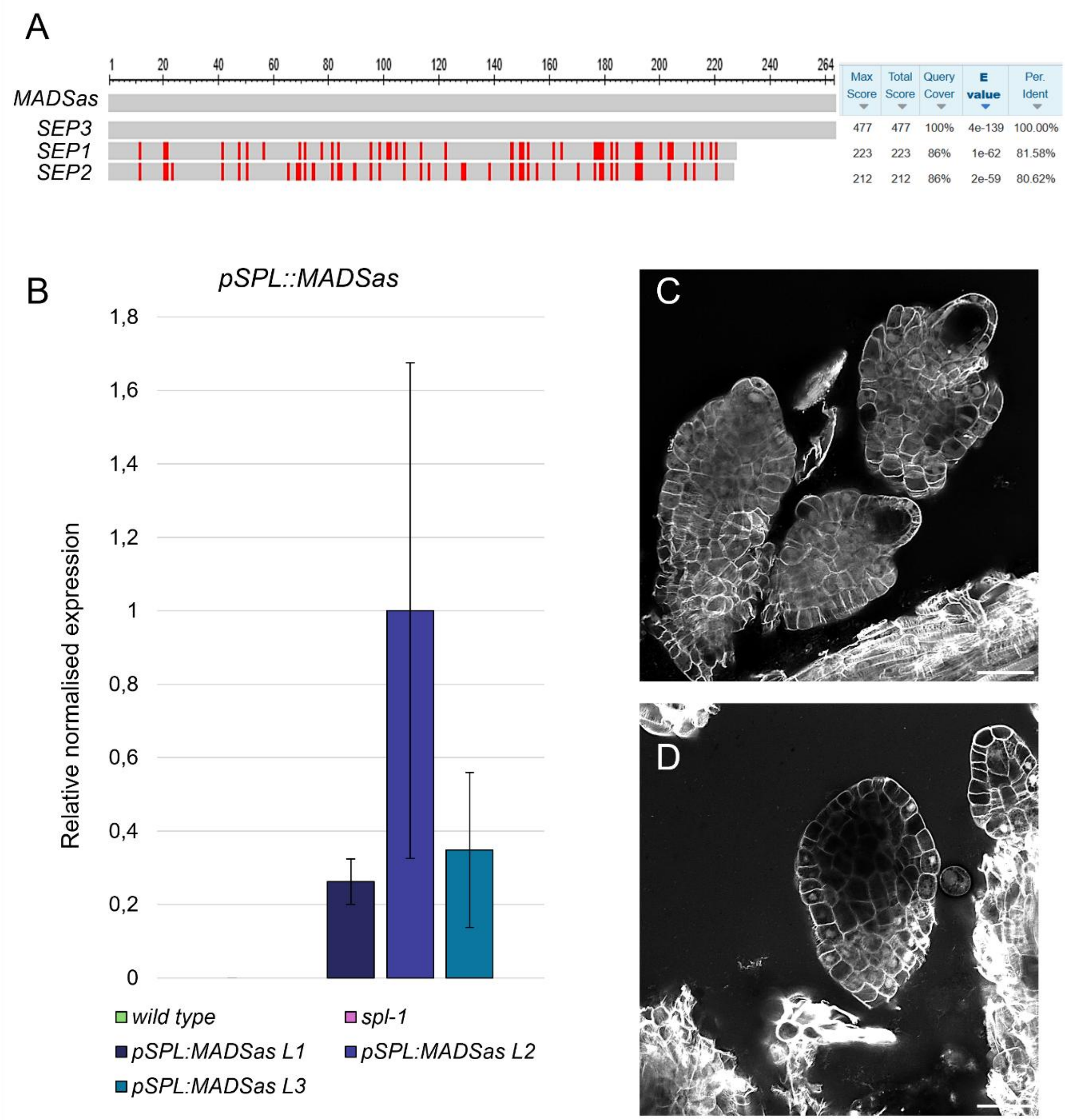


**Supplementary Figure 5: *MADSas* sequence homology to *SEP1, 2* and 3 and *MADSas* expression and ovule phenotypes in *pSPL/NZZ::MADSas* lines.** (**A**) Multiple sequence alignment among *MADSas, SEP1, SEP2* and *SEP3* transcripts performed with the NCBI BLASTn tool. The *MADSas* is transcribed from a 264bp portion amplified from *SEP3 MADS* box sequence and placed in reverse under the control of the *SPL/NZZ* promoter. The *MADSas* has around 80% of sequence homology with *SEP1* and *2.* In the schematic of the alignment, mismatched bases are depicted in red. (**B**) Bar plot showing *pSPL/NZZ::MADSas* relative normalised expression in inflorescences from wild type, *spl-1* and the three independent *pSPL/NZZ::MADSas* lines. *ACTIN8* was used as the housekeeping gene. Bars represent the mean ± SEM of the expression, as evaluated from three technical replicates per each sample. Primers used are listed in Supplementary Data 4. (**C**, **D**)

Examples of ovule phenotypes observed in *pSPL/NZZ::MADSas* lines. While a portion of the ovule did not develop the MMC (**C**), the structure of few ovules appears severely impaired, hindering the identification of the different ovule domains (**D**). Such ovules (**D**) were not considered for the evaluation of the MMC differentiation. Scale Bars: 20 µm.

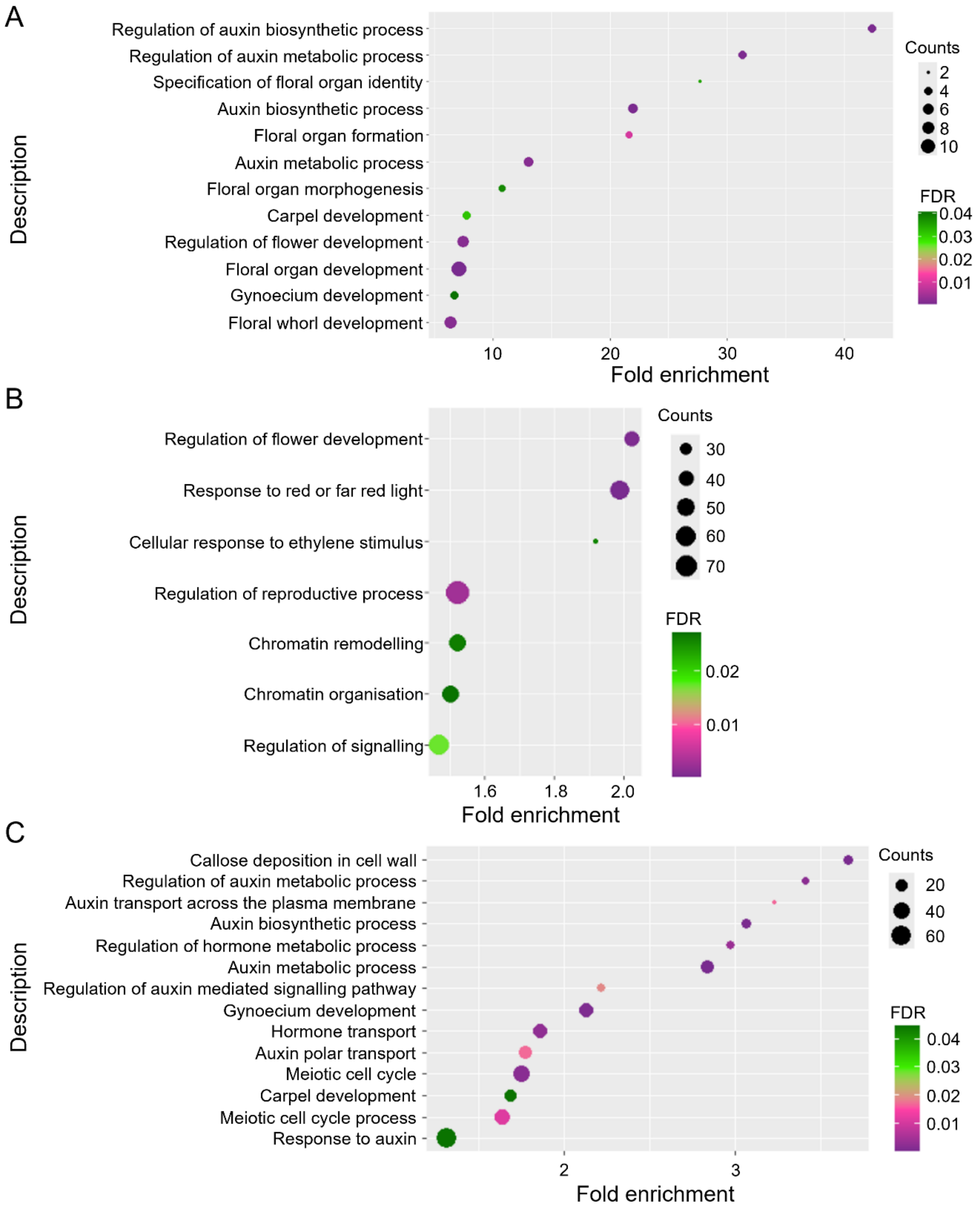


**Supplementary Figure 6: Biological processes GO terms enrichment analysis of SPL/NZZ and SEP3 common target genes and of DE genes in the *spl-1* pistil with respect to the wild type.** (**A**) Biological process GO term enrichment analysis, done on the 185 genes that have overlapping peaks between SPL/NZZ and SEP3 ChIPseq experiments. Categories associated to auxin and hormone processes, flower and floral organs development are enriched. Enriched GO terms with an FDR ≤ 0.05 were considered significant.

(**B**) Biological process GO terms enrichment analysis conducted on the downregulated genes in *spl-1* pistils with respect to wild type. Categories associated to flower development and chromatin organisation are among the most enriched categories. (**C**) Biological process GO terms enrichment analysis conducted on the upregulated genes in *spl-1* pistils with respect to wild type. Categories associated to meiosis, gynoecium development, auxin and hormones processes are among the most enriched categories (Supplementary Data 3).

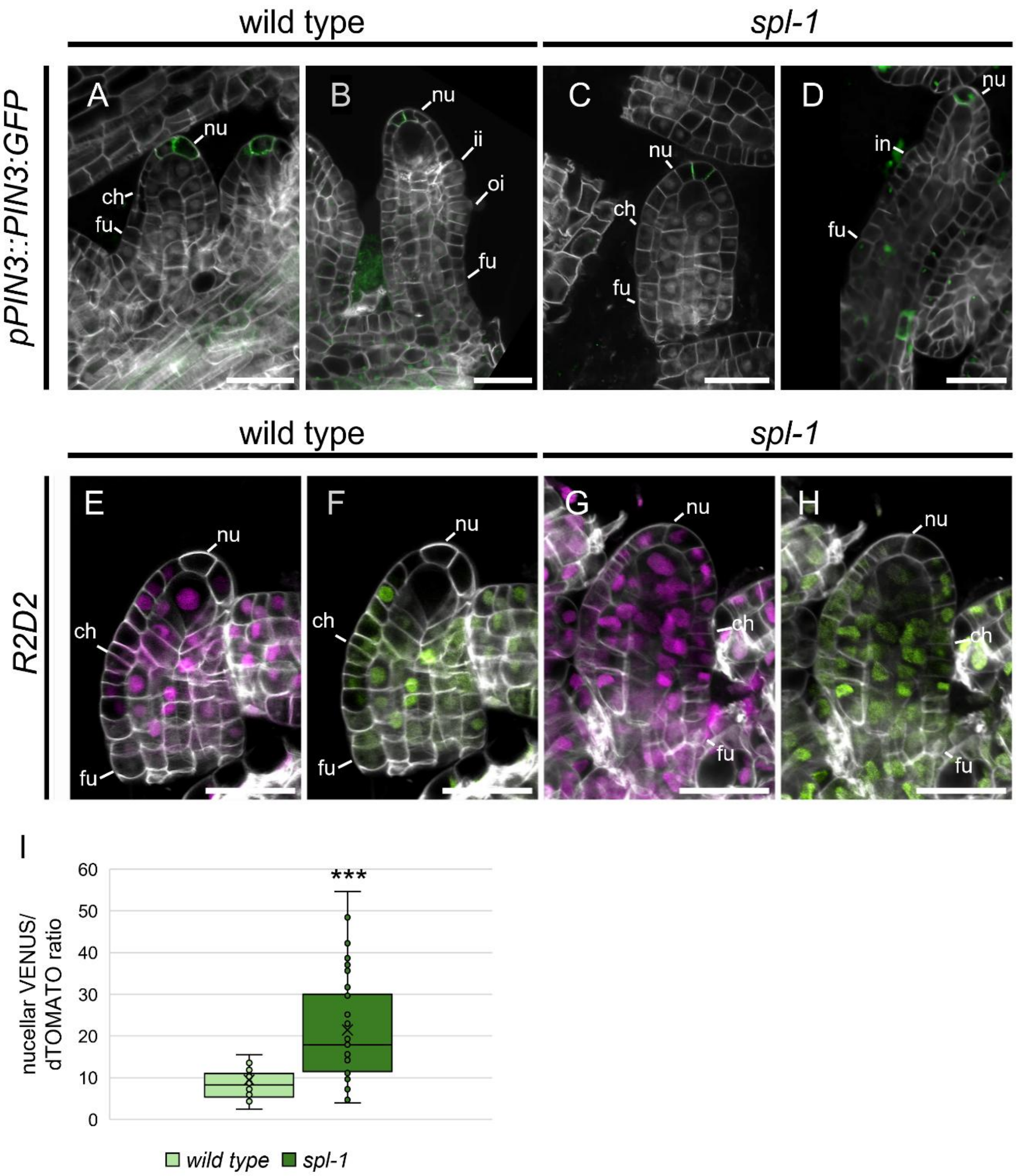


**Supplementary Figure 7: *pPIN3::PIN3:GFP* expression in wild type and *spl-1* ovules and *R2D2* signal in wild type and *spl-1* ovules at stage 1-II.** (**A, B**) PIN3-GFP accumulation at stage 2-I (**A**) and 2-II (**B**) in wild type. (**C, D**) PIN3-GFP accumulation at stage 2-I (**C**) and 2-II (**D**) in *spl-1*. (**E-H**) *R2D2* reporter line in wild type (**E**, **F**) and *spl-1* (**G**, **H**) ovules at stage 2-I, showing mDII-tdTOMATO (**E**, **G**) and DII-VENUS (**F**, **H**) accumulation. (**I**) Box plot showing the tdTOMATO/VENUS signal ratio from the *R2D2* reporter, in the wild type and *spl-1* nucella at stage 2-III. The analysis has been performed on 28 and 45 nuclei, from 3 wild type and 4 *spl-1* different ovules, respectively. Asterisks over boxes represent the statistical significance as determined by student’s t-test, two-tailed distribution homoscedastic, confronting the mutant with the wild-type condition. *** = $p < 0.001$. Box-plots elements correspond to: centre line = median; X = box limits = interquartile range; whiskers = lowest and highest values in the 1.5 interquartile range. Single measures are represented with dots in the boxes. Abbreviations: nu= nucellus; ch= chalaza; fu= funiculus; ii= inner integument; oi= outer integument; in= integument. Scale bar = 20 µm.

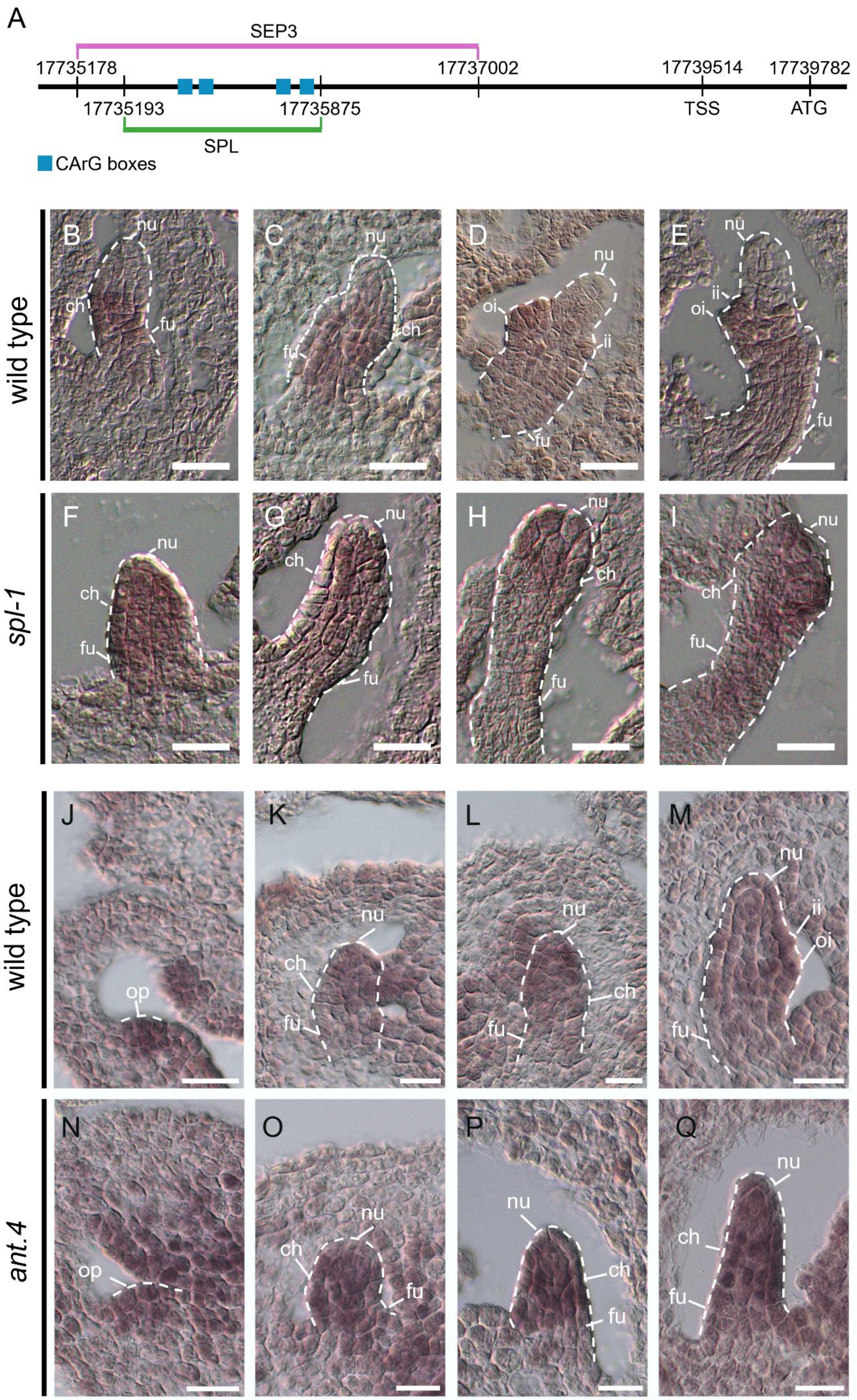
A
SEP3
17735178
17737002
17739514
17739782
17735193
17735875
TSS
ATG
SPL
CArG boxes
wild type
spl-1
wild type
ant.4
B C D E F G H I J K L M N O P Q
nu
ch
fu
oi
ii
op

**Supplementary Figure 8: SPL/NZZ and SEP3 ChIPseq peaks on *ANT* promoter, *ANT* expression in wild type and *spl-1* ovules and *PIN1* expression in wild type and *ant.4* ovules.** (**A**) Schematic of the *ANT* promoter region showing, with a green box, the peak identified in the SPL/NZZ ChIPseq and in magenta the peak identified in the SEP3 ChIPseq. The 4 CArG-box sequences identified by the MEME analysis on the SPL/NZZ peak are highlighted by blue boxes. (**B**-**I**) *ANT* transcript *in-situ* hybridisation on wild type (**B-E**) and *spl-1* (**F**-**I**) ovules. In wild-type ovules, *ANT* is expressed in the chalaza and the funiculus (**B-E**) whereas, *ANT* transcript accumulates ectopically also in the nucellus in the *spl-1* mutant (**F-I**). (**J**-**Q**) *PIN1* expression detected by *in-situ* hybridisation in wild type (**J**-**M**) and *ant.4* (**N**-**Q**) ovules at stages 0-II (**J**, **N**), 1-II (**K**, **O**), 2-I (**L**, **P**) and 2-III (**M**, **Q**). At stages 2-I, *PIN1* expression could be observed mainly in the nucellus of the wild-type ovule (**L**). At stage 2-III (**M**), *PIN1* is still visible in the nucellus and partially in the chalaza. By contrast, in the *ant.4* ovule (**N**-**Q**), *PIN1* expression can be observed in the whole ovule primordia. Abbreviations: op= ovule primordia; nu= nucellus; ch= chalaza; fu= funiculus; ii= inner integument; oi= outer integument. Scale bars = 20µm.

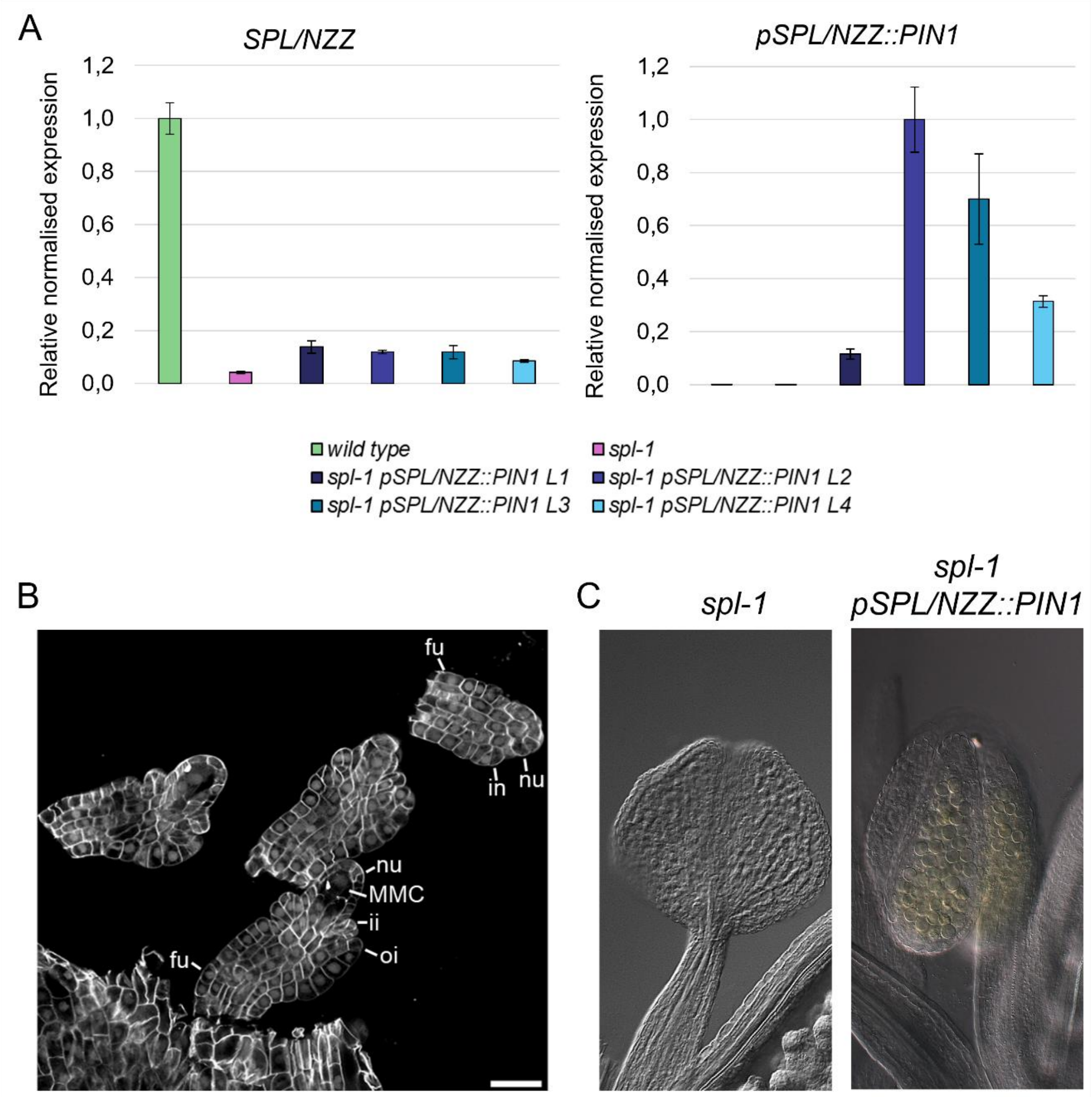


**Supplementary Figure 9: *pSPL/NZZ::PIN1* expression and ovule and anther phenotypes in *spl-1 pSPL/NZZ::PIN1* plants.** (**A**) Bar plot showing *SPL/NZZ* relative normalised expression in inflorescences from the wild type, *spl-1* and four independent *spl-1 pSPL/NZZ::PIN1* lines. (**B**) Bar plot showing *pSPL/NZZ::PIN1* relative normalised expression in the same samples presented in (**A**). *ACTIN8* was used as the housekeeping gene. Bars represent the mean ± SEM of the expression, as evaluated from three technical replicates per each sample. Primers used are listed in Supplementary Data 4. (**C**) Despite the ability of *pSPL/NZZ::PIN1* to rescue the MMC specification, few ovules still resemble the *spl-1* phenotype. (**D**) *spl-1* and *spl-1 pSPL/NZZ::PIN1* anthers. In contrast to the *spl-1* situation, *spl-1 pSPL/NZZ::PIN1* could develop anthers generating wild-type-like pollen grains. Scale bar = 20 μm.

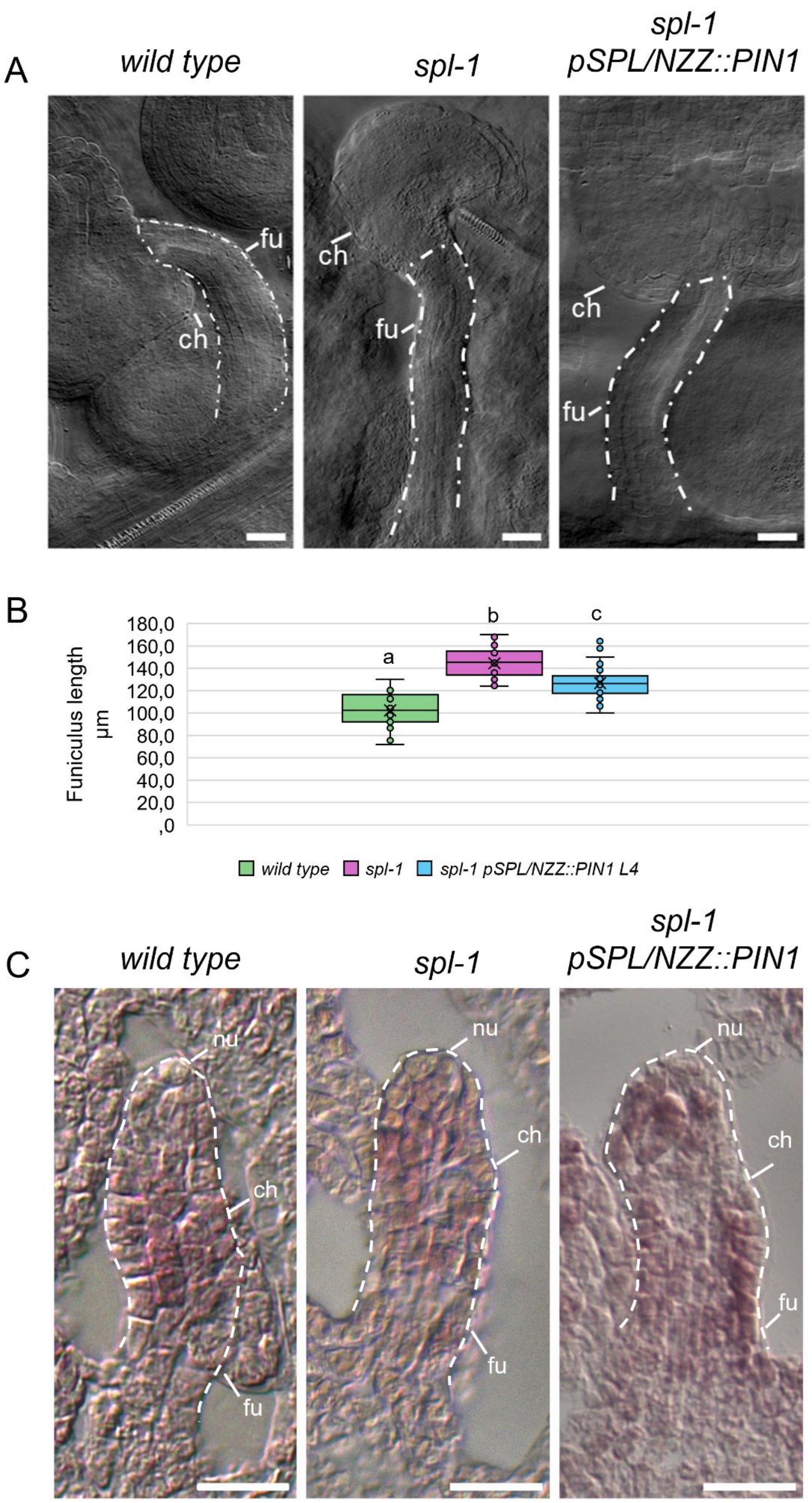
A
wild type
spl-1
spl-1 pSPL/NZZ::PIN1
fu
ch
B
Funiculus length µm
180,0
160,0
140,0
120,0
100,0
80,0
60,0
40,0
20,0
,0
a
b
c
wild type
spl-1
spl-1 pSPL/NZZ::PIN1 L4
C
nu
ch
fu

**Supplementary Figure 10: Funiculus length and *ANT* expression in *spl-1 pSPL/NZZ::PIN1.*** (**A**, **B**) Funiculus length in wild type, *spl-1* and *spl-1 pSPL/NZZ::PIN1* ovules at stage 3-VI. Even though the ovule development was restored in *spl-1 pSPL/NZZ::PIN1* (**A**)*,* funiculi were still statistically longer than the wild-type ones. Nevertheless, *spl-1 pSPL/NZZ::PIN1* funiculi resulted statistically shorter than the *spl-1* ones (**B**). Measurements were performed on 25, 33 and 45 different funiculi, respectively, for the wild type, *spl-1* and *spl-1 pSPL/NZZ::PIN1 L4.* Letters above the box plots indicate homogenous categories with $p < 0.05$, as determined by one-way ANOVA with post-hoc Tukey HSD test. Box-plots elements correspond to: centre line = median; X= average; box limits = interquartile range; whiskers = lowest and highest values in the 1.5 interquartile range. Single measures are represented with dots in the boxes. (**C**) *ANT* transcript *in-situ* hybridisation on wild type, *spl-1* and *spl-1 pSPL/NZZ::PIN1 L4* ovules. Despite the MMC differentiation is restored in *spl-1 pSPL/NZZ::PIN1,* SPL direct targets as *ANT* remains ectopically expressed, similarly to the *spl-1* situation. Abbreviations: nu= nucellus; ch= chalaza; fu= funiculus. Scale bars = 20 µm.

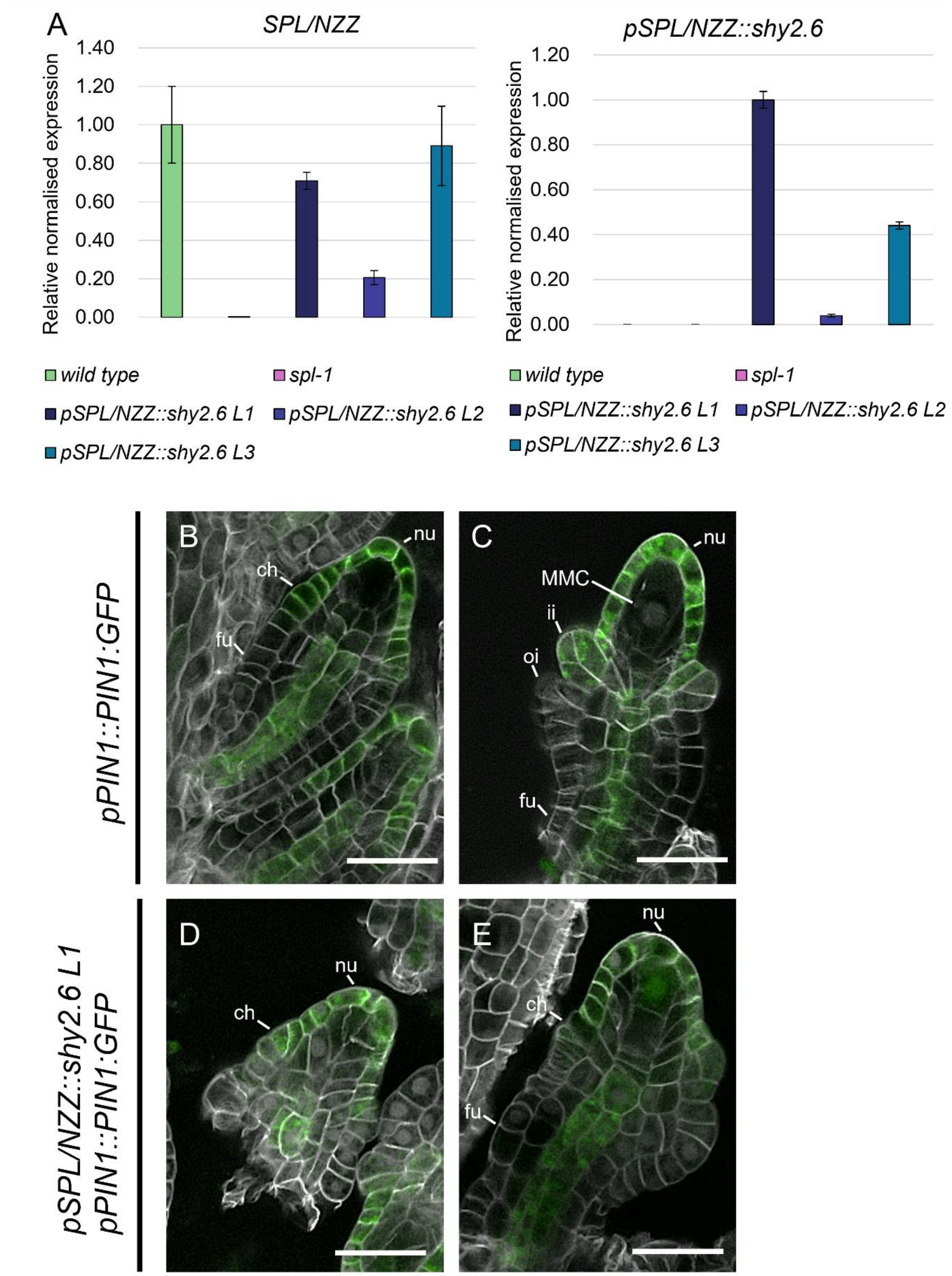


**Supplementary Figure 11: *pSPL/NZZ::shy2.6* expression level and PIN1-GFP accumulation in *pSPL/NZZ::shy2.6.*** (**A**) Bar plot showing *SPL/NZZ* and *pSPL/NZZ::shy2.6* relative normalised expressions in inflorescences from wild type, *spl-1* and three independent *pSPL/NZZ::shy2.6* lines. *ACTIN8* was used as housekeeping gene for normalisation. Bars represent the mean ± SEM of the expression, as evaluated from three technical replicates per each sample. Primers used are listed in Supplementary Data 4. (**B**-**E**) PIN1-GFP accumulation in wild type (**B**, **C**) and *pSPL/NZZ::shy2.6 L1* (**D**, **E**) ovules at stages 2-I (**B**, **D**) and 2-III (**C**, **E**). Abbreviations: nu= nucellus; ch= chalaza; fu= funiculus; ii= inner integument; oi= outer integument; MMC= Megaspore mother cell. Scale bar = 20 μm.

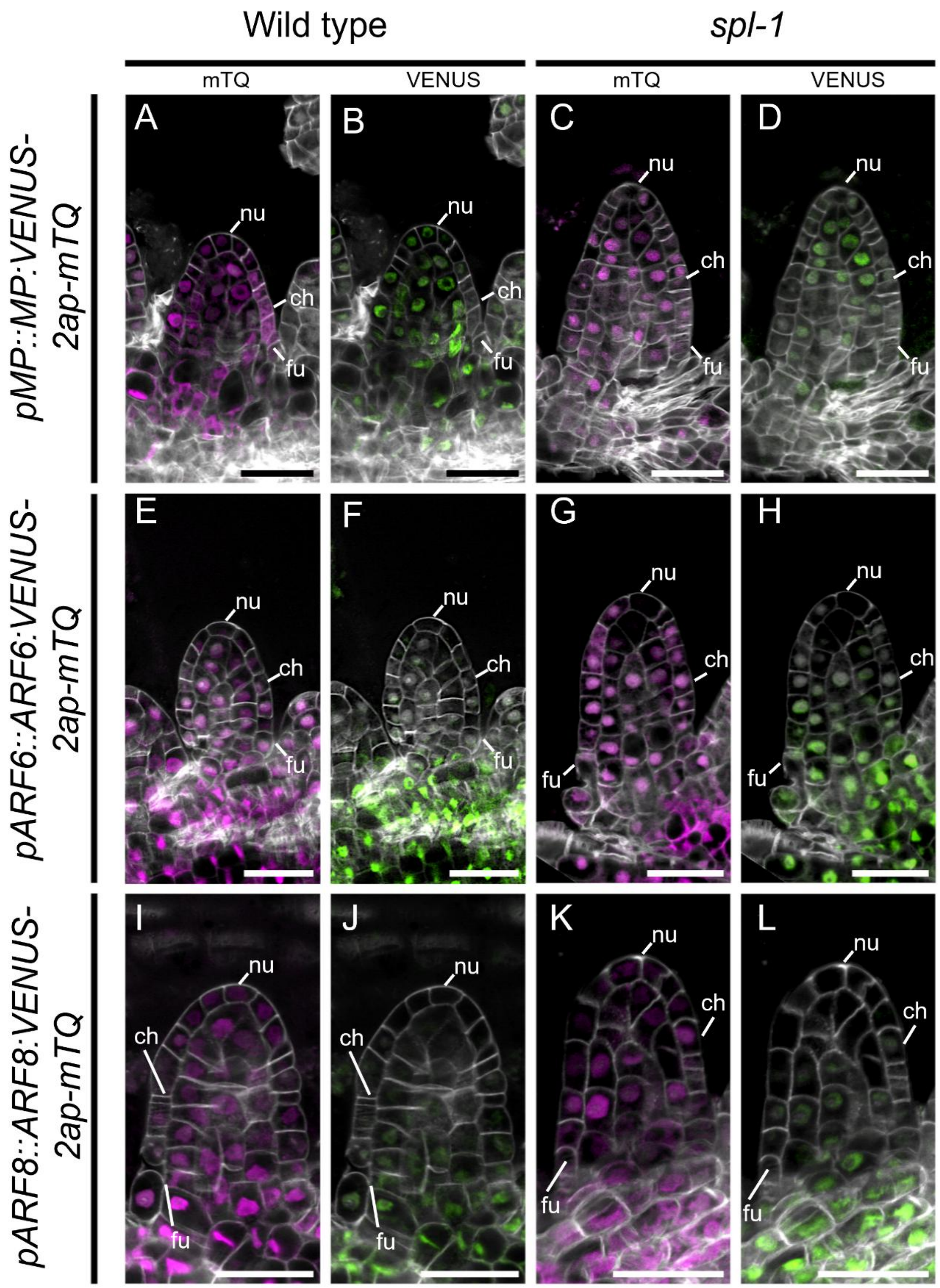


**Supplementary Figure 12: ClassA ARFs expression in wild type and *spl-1* ovules.** (**A-D**) *pMP::MP:VENUS-2ap-mTQ* reporter in wild type (**A**, **B**) and *spl-1* (**C**, **D**) ovules. (**E-H**) *pARF6::ARF6:VENUS-2ap-mTQ* reporter in wild type (**E**, **F**) and *spl-1* (**G**, **H**) ovules. (**I-L**) *pARF8::ARF8:VENUS-2ap-mTQ* reporter in wild type (**I**, **J**) and *spl-1* (**K**, **L**) ovules. *MP, ARF6* and *ARF8* domains of expression and translation are visualised thanks to the mTQ signal (**A**, **C**, **E**, **G**, **I**, **K**), while domains of protein accumulation are visualised by the VENUS signal (**B**, **D**, **F**, **H**, **J**, **L**). Abbreviations: nu= nucellus; ch= chalaza; fu= funiculus. Scale bar = 20 µm.

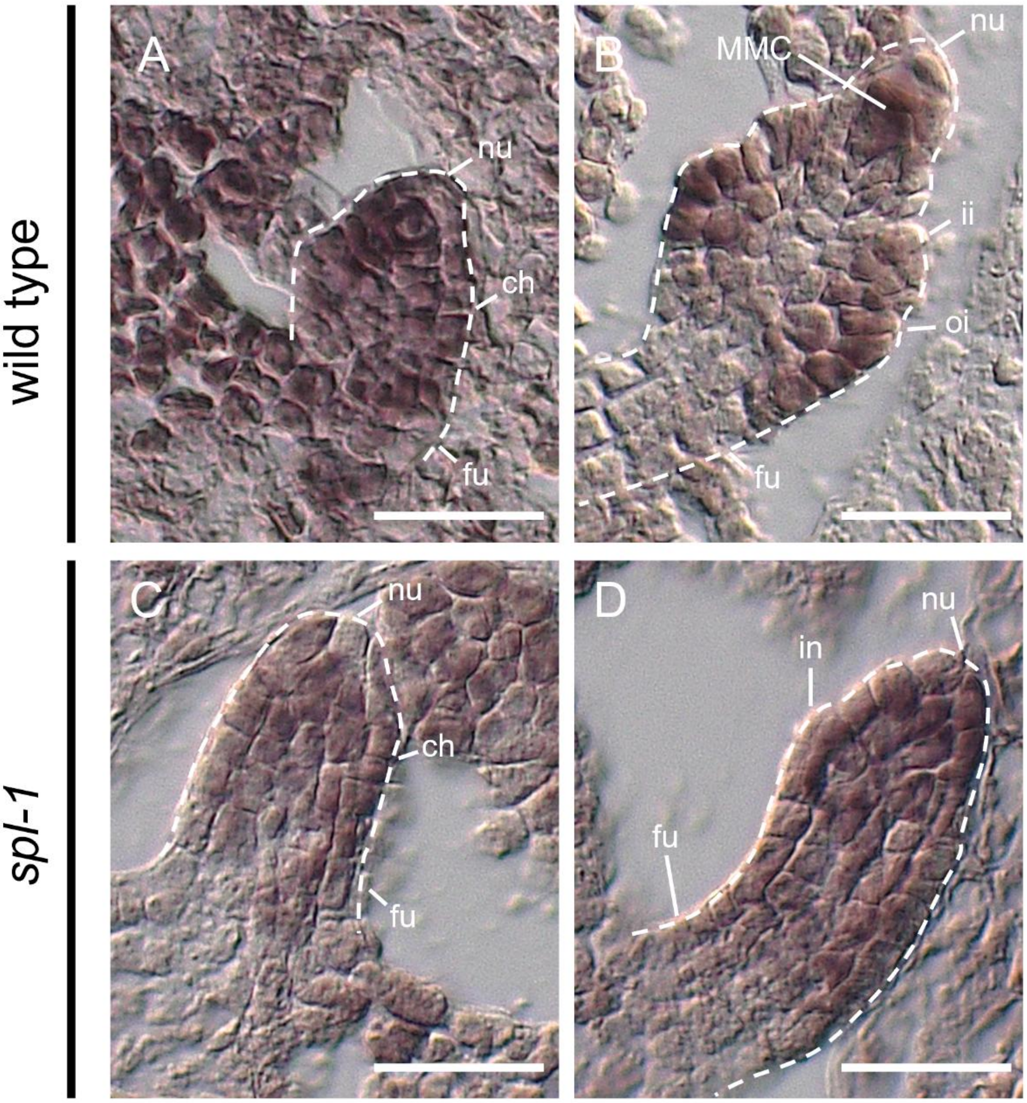


**Supplementary Figure 13: ARF9 expression in wild type and *spl-1* ovules detected by *in-situ* hybridisation.** (**A, D**) ISH showing *ARF9* expression in wild type ovules (**A**, **B**) and *spl-1* (**C, D**) ovules at stages 1-II (**A, C**) and 2-III (**B, D**). Abbreviations: nu= nucellus; ch= chalaza; fu= funiculus; MMC= megaspore mother cell; ii= inner integument; oi= outer integument; in= integument. Scale bar = 20 µm.

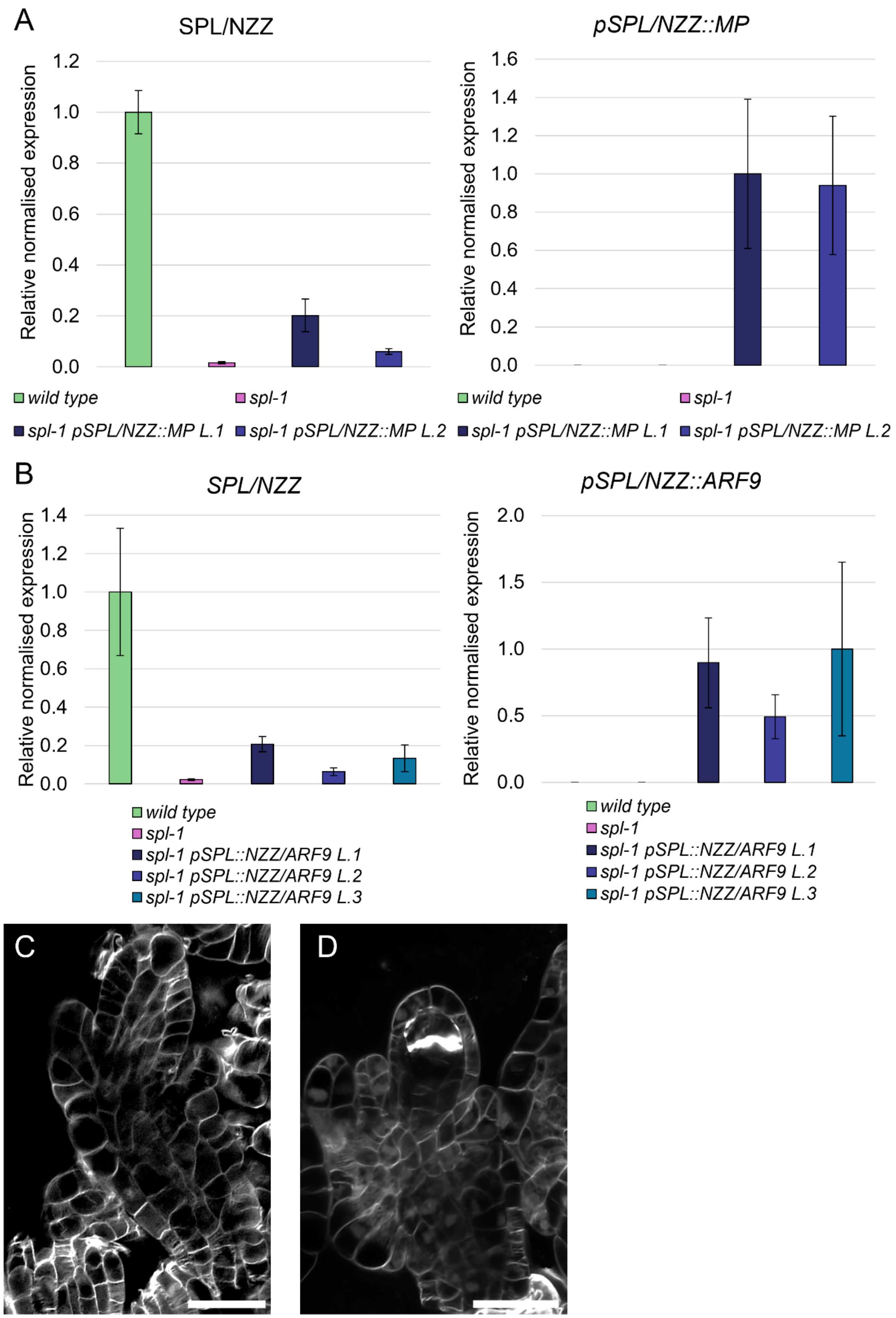
A
SPL/NZZ
Relative normalised expression
1.2
1.0
0.8
0.6
0.4
0.2
0.0
pSPL/NZZ::MP
1.6
1.4
wild type
spl-1
spl-1 pSPL/NZZ::MP L.1
spl-1 pSPL/NZZ::MP L.2
B
SPL/NZZ
pSPL/NZZ::ARF9
2.0
1.5
0.5
spl-1 pSPL::NZZ/ARF9 L.1
spl-1 pSPL::NZZ/ARF9 L.2
spl-1 pSPL::NZZ/ARF9 L.3
C
D

310

311

**Supplementary Figure 14: *pSPL/NZZ::MP* and *pSPL/NZZ::ARF9* expression, *spl-1 pSPL/NZZ::MP* and *spl-1 pSPL/NZZ::ARF9* ovules at stage 2-IV.** (**A**) Bar plot showing *SPL/NZZ* and *pSPL/NZZ::MP* relative normalised expressions in inflorescences from wild type, *spl-1* and two independent *spl-1 pSPL/NZZ::MP* lines. *ACTIN8* was used as housekeeping gene for normalisation. Bars represent the mean ± SEM of the expression, as evaluated from three technical replicates per each sample. Primers used are listed in Supplementary Data 4. (**B**) Bar plot showing *SPL/NZZ* and *pSPL/NZZ::ARF9* relative normalised expressions in inflorescences from wild type, *spl-1* and three independent *spl-1 pSPL/NZZ::ARF9* lines. *ACTIN8* was used as housekeeping gene for normalisation. Bars represent the mean ± SEM of the expression, as evaluated from three technical replicates per each sample. Primers used are listed in Supplementary Data 4. (**C**) In *spl-1 pSPL/NZZ::MP* ovules, the MMC does not proceed into the meiotic process. (**D**) In *spl-1 pSPL/NZZ::ARF9,* the MMC divide meiotically, as indicated by the thik callose deposition associated with the end of the meiosis I. Scale bars: 20 μm.

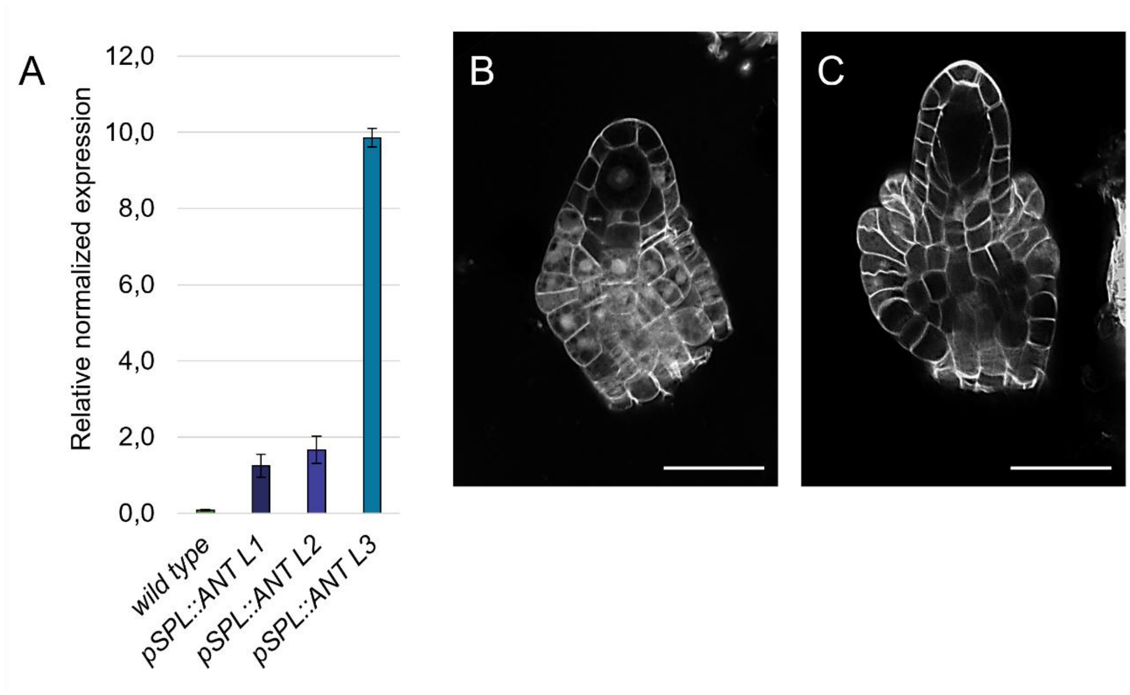


**Supplementary Figure 15: Expression of *pSPL/NZZ::ANT* and ovules phenotypes in *pSPL/NZZ::ANT* lines.** (**A**) Bar plot showing and *pSPL/NZZ::ANT* relative normalised expressions in inflorescences from wild type and three independent *pSPL/NZZ::ANT* lines. *ACTIN8* was used as housekeeping gene for normalisation. Bars represent the mean ± SEM of the expression, as evaluated from three technical replicates per each sample. Primers used are listed in Supplementary Data 4. (**B**, **C**) Ovule development in *pSPL/NZZ::ANT* lines. Scale bars: 20 μm.

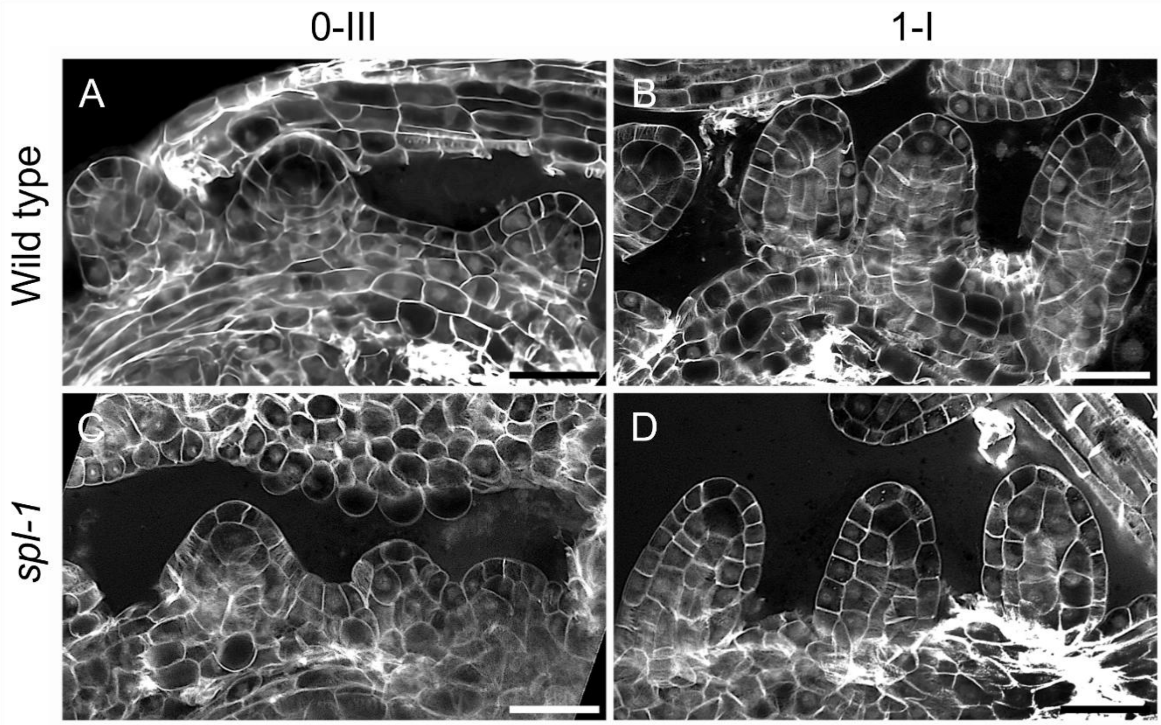


**Supplementary Figure 16: Wild type and *spl-1* ovules at precocious developmental stages.** (**A, B**) wild type ovules at stages 0-**III** (**A**) and 1-**I** (**B**). (**C**, **D**) *spl-1* ovules at stages 0-**III** (**C**) and 1-**I** (**D**). Before MMC specification, wild type and *spl-1* share similar phenotype, showing the presence of putative MMC precursors in the nucellus. Scale bar: 20 µm.
